# Optimal Individualized Combination Immunotherapy/Oncolytic Virotherapy Determined Through *In Silico* Clinical Trials Improves Late Stage Melanoma Patient Outcomes

**DOI:** 10.1101/585711

**Authors:** Tyler Cassidy, Morgan Craig

## Abstract

Oncolytic virothcrapics, including the modified herpes simplex virus talimogene laherparepvec (T-VEC), have shown great promise as potent instigators of anti-tumour immune effects. The OPTiM trial in particular demonstrated the superior anti-cancer effects of T-VEC as compared to more traditional immunotherapy treatment using exogenous administration of granulocyte-macrophage colony-stimulating factor (GM-CSF). Theoretically, a combined approach leveraging immunotherapies: like exogenous cytokine administration and oncolytic virotherapy would elicit an even greater immune response and improve patient outcomes, but given that their efficacy and safety must be tested in large clinical trials, combination therapeutic regimens have yet to be established. By adopting computational biology and *in silico* clinical trial approaches, here we show significantly improved patient outcomes for individuals with late-stage melanoma by personalizing and optimizing combination oncolytic, virotherapv and GM-CSF therapy. Our results serve as a proof-of-concept, for interdisciplinary approaches to determining combination therapy, and suggest promising avenues of investigation towards tailored combination immunotherapy/oncolytic virotherapy.

## Introduction

Modern cancer treatments increasingly incorporate a broad class of biological therapies known as immunotherapies to activate the immune system against cancer cells in a generalized or targeted way.^1,2^ These therapies seek to exploit existing tumour-immune interactions to more efficiently and effectively recognize and destroy tumour cells while hopefully minimizing off-target and detrimental side effects. Current and investigational immunotherapies include immune-checkpoint inhibitors, monoclonal antibodies, CAR-T cells, and the exogenous administration of cytokines. One such cytokine, granulocyte-macrophage colony-stimulating factor (GM-CSF), is a white blood cell growth factor responsible for stimulating granulocyte production, and orchestrating innate inflammatory responses. GM-CSF has been used to increase the efficacy of monoclonal antibodies, and has also been administered during B-cell lymphoma treatment to activate certain immune cell, subsets.^2^

Another older idea, more recently explored, is to use oncolytic viruses to destroy tumour cells^3,4^ and activate an immune response. These oncolytic viruses are genetically engineered to preferentially attack and infect cancerous cells,^5,6^ forcing them to undergo lysis and release tumour specific antigens that signal the immune system to mount an anti-tumour response.^7,8^ This double effect, against tumour cells has motivated the study of oncolytic viruses as a treatment against a variety of malignant solid tumours, and the modified herpes simplex virus talimogene laherparepvec (T-VEC) was approved for use in patients with non-resectable melanoma in 2015.^9-11^ T-VEC is specifically engineered to enhance expression of GM-CSF after viral infection into tumour cells.^9^ However, despite much promise, the efficacy of oncolytic virus monotherapy has been limited.^8,12,13^ As it is reasonable to expect that immunotherapy and virotherapy could act synergistically to instigate an immune response against tumour cells,^14-16^ recent efforts have focused on determining the anticipated benefit to their use in combination with a variety of immunotherapies.^17,18^ To that end, GM-CSF has been considered as an immune stimulant during oncolytic virotherapy.^2^

Generally speaking, combination therapy can carry a high therapy burden and may increase overall toxicity. ^13,18^ Unfortunately, running clinical trials for all possible (dose, time)-pairs of a proposed combined treatment to determine efficient and safe scheduling is both time and cost prohibitive. As a consequence, understanding the regimen scheduling of combination immuno-/oncolytic virotherapy remains an open problem. Fortunately, computational biology and *in silico* predictions provide a platform for the non-invasive exploration of potential therapeutic schedules that concretely improve patient outcomes.^19-21^ To that end, here we detail a rational, quantitative approach to therapy scheduling and optimization of combination immuno-oncolytic virotherapy for patients with late stage melanoma using an *in silico* clinical trial.

By generating identical virtual patient cohorts, we determined optimal, individualized treatment regimens for combined GM-CSF immunotherapy and oncolytic virotherapy through *in silico* clinical trial simulations. These individualized schedules served to determine an optimal dosing scheme that significantly improves overall survival while substantially reducing drug burden, highlighting the potential and potency of rational regimen prediction using a computational biology approach.

## Materials and Methods

### Computational Biology Model

To investigate the synergy between immunotherapy (exogenous GM-CSF) and oncolytic virotherapy, we adapted our previous mathematical model^22^ describing the instantaneous change in tumour size, phagocyte numbers and cytokine concentrations over time, with and without therapy. The model tracks both immuno-susceptible and immuno-resistant tumour cell populations as they transit through the cell cycle. Quiescent immuno-susceptible tumour cells can be cleared through either random death or immune pressure, or transit into the *G*_1_ phase to begin division. Cells in *G*_1_ are also subject to random death and immune clearance before beginning the mitotic process. After completing division, susceptible cells return to quiescence. Mitotic cells may mutate at rate *µ* into an immuno-resistant cell type with a low probability. This immuno-resistant lineage maintains the same cell cycle behaviour of non-resistant tumour cells, but, as it evades immune pressure, is not subject to any immune interactions. We do not distinguish between different types of immune cells in the tumour microenvironment, but rather model all phagocytes as a single population. These phagocytes interact with the susceptible tumour cell population and produce a pro-inflammatory cytokine (e.g. interleukin-12, tumour necrosis factor, interferon gamma, GM-CSF etc.) to recruit other phagocytes to the tumour site. Model predictions are obtained by simulation, as previously described^22^ (full details are provided in the Supplementary Information). The various interactions described above are schematized in Figure 1.

**Figure 1:**
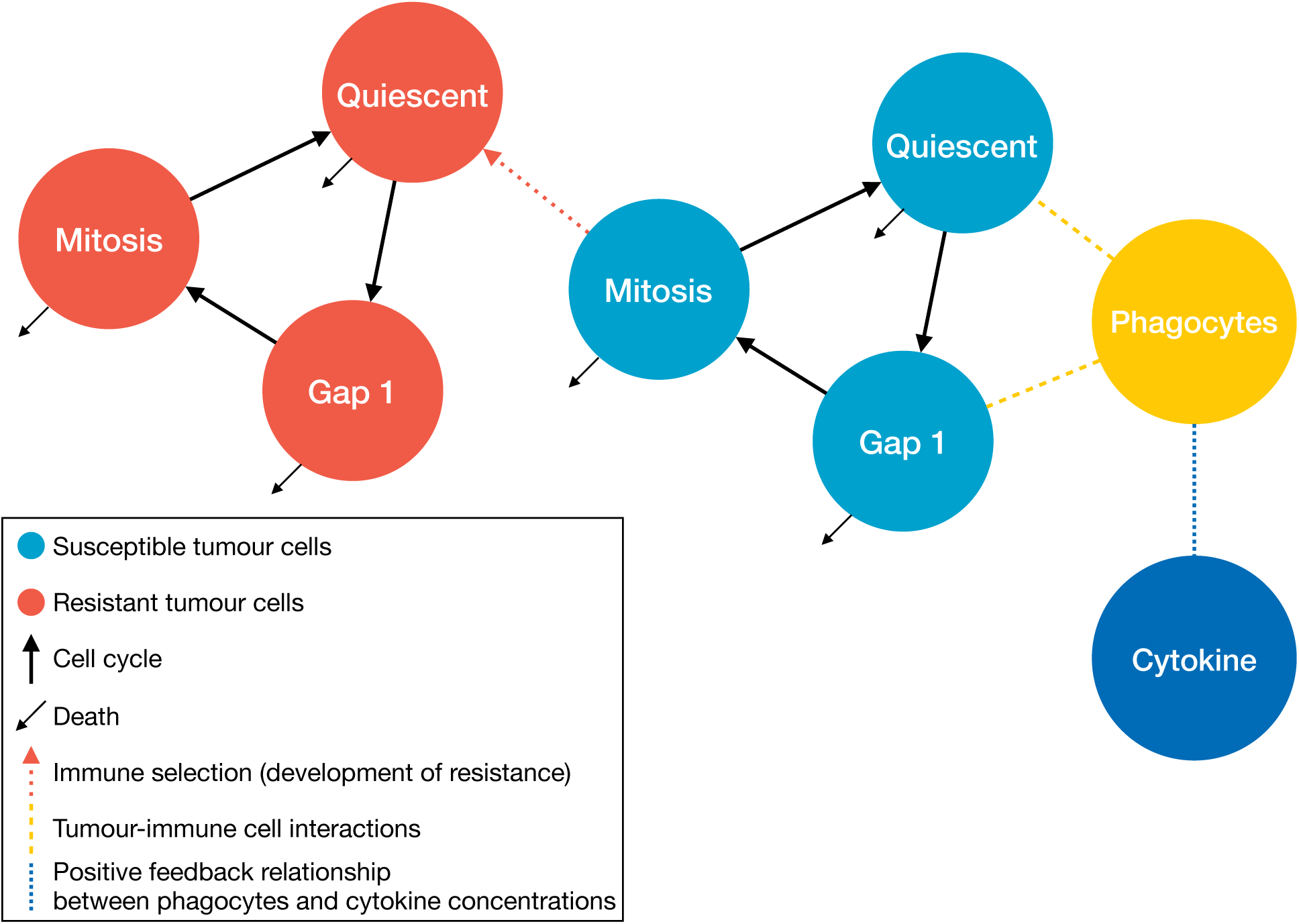
Pictorial representation of the tumour growth model. Quiescent cells activate to begin division by transiting into the *G*_*1*_ phase of the cell-cycle. Cells exit *G*_1_ to enter the active phase and complete division. Most susceptible cells in the active phase re-enter quiescence after mitosis, however certain dividing cells may mutate into an immuno-resistant. lineage (red dotted arrow). Immune interactions are driven by phagocytes who come into contact with quiescent and *G*_1_ phase susceptible cells (dashed yellow lines). Tumour-immune interactions increase pro-inflammatory cytokine concentrations to recruit additional phagocytes to the tumour site (blue dotted line). Cells and cytokine are denoted by circles, processes by squares, and rates by arrows.

### Generation of *In Silico* Individuals and Patient Cohorts

In our previous work, the growth of a malignant tumour was parameterized using average values representative of an average patient.^22^ However, as patient populations are heterogenous, our mathematical model must be adapted to reflect this reality. Accordingly, we individualized the model by generating a unique set of parameters to represent individuals as potential participants in the virtual trial. We sampled from a generated normal distribution with mean as previously given^22^ and with realistic standard deviations about this mean. Mathematically, we considered **p** to be the vector of average parameter values from our previous model.^22^ Each individual is then generated by sampling from a normal distribution with mean *µ* = **p** and standard deviation *σ* = 16.33%, so that 99.7% of values fall within the interval [*µ* –3*σ, µ* + 3*σ*] = [0.5**p**, 1.5**p**].

To mimic “real” clinical trials, we imposed selection and inclusion criteria on each generated individual by verifying that each patient responds in a physiologically-realistic way without and with treatment.^19^ For this, we compared the predicted response of each virtual patient to currently approved oncolytic virotherapy for stage IIIb or IV non-surgically resectable melanoma.^8,9^ Specifically, we assessed whether the predicted tumour doubling time of each individual corresponded to clinically relevant tumour doubling times.^23^ Tumour doubling time therefore represented the sole inclusion criterion for subsequent, enrolment, *in silico* clinical trial simulations.

We accepted a total of 200 virtual patients, generated by the parameter sampling and selection processes outlined above. Each virtual patient was then reproduced into *n* identical clones, and each resulting clone was subsequently assigned to one of *n* separate cohorts (for example, a treatment free control group, a mono-immunotherapy group, and an oncolytic virotherapy group, for a total of *n* = 3 cohorts). In this way, the total number of participants is 200 times the total number of simulated investigational arms, or 200*n*. The *in silico* trial generation process is schematized in Figure 2.

**Figure 2:**
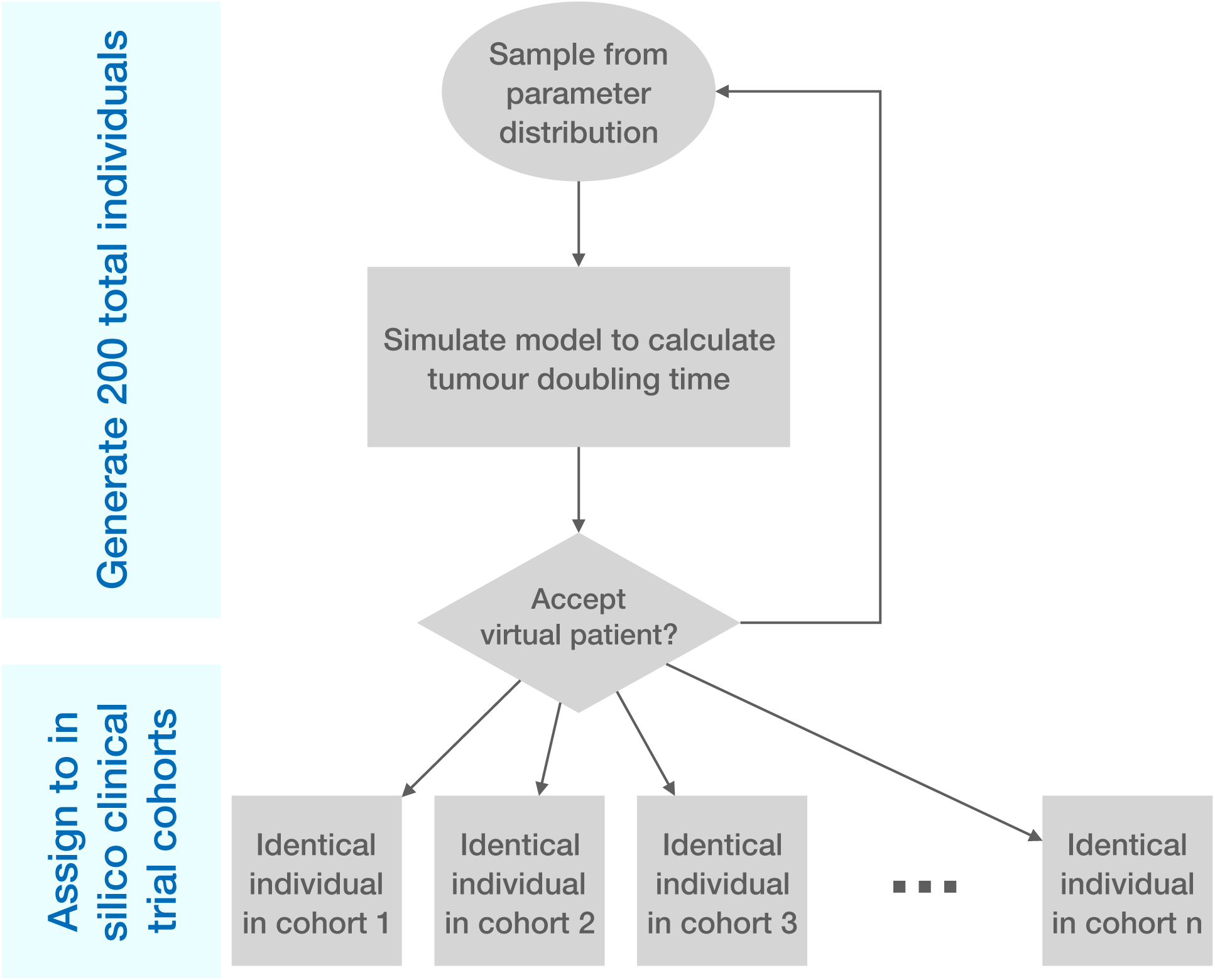
A pictoral representation of the *in silico* patient creation algorithm. Individual *in silico* patient parameter values are sampled from a normal distribution of values based on an average parameterization^22^. The model was then simulated for each individual and tumour growth predictions (tumour doubling time) were tested for physiological relevance. If the *in silico* patient’s tumour growth behaviour was considered physiologically realistic, they were cloned n times and each clone was assigned to *n* separate arms of the *in silico* clinical trial.

### Recapitulation of Previous Trial Data

Using three identical cohorts, we evaluated patient outcomes when they received no treatment (Cohort 1), immunotherapy (Cohort 2), or oncolytic virus monotherapy (Cohort 3). This setup mimics that of the T-VEC OPTiM trial, where individuals were randomized to receive either intralesional T-VEC or subcutaneous GM-CSF.^9^

In both the *in silico* immunotherapy and oncolytic virus monotherapy cases, the dosing schedules were identical to the ones used in the OPTiM trial:^9^ patients in the T-VEC arm received a priming dose of 10^6^ plaque forming units pfu/mL, followed by 10^8^ pfu/mL doses to a maximal total administration of 4 mL per treatment. T-VEC was administered every 14 days Patients in the GM-CSF arm received 125 *µ*g/m^2^ of subcutaneous GM-CSF administered on 14 consecutive days followed by 14 days of no treatment. In both arms, treatment continued for up to 12 months but could be discontinued due to disease progression, intolerability, or the disappearance of injectable lesions.

We fixed an oncolytic virotherapy dose of 1 x 10^6^ virions, as the amount of virus administered the original trial varied based on both patients and physicians.^9^ Note that the units between the OPTiM trial and our *in silico* trial differ owing only to the units of the mathematical model’s parameters and the conversion of pfu to infectious virions. Individuals receiving GM-CSF immunotherapy in the *in silico* trial were administered 125 *µ*g/m^2^ of GM-CSF daily for 14 days in 28 day cycles. For both arms, we simulated the model over a fixed treatment time of 12 months.

Late stage melanoma has a low survival rate.^24^ Mortality as a function of tumour doublings has been estimated to occur between 40 and 45 tumour doublings.^23,25,26^ Given that roughly 30 doublings occur before clinical presentation,^26^ we estimated that there are approximately 10 and 15 tumour doublings between diagnosis and death. *In silico* patients were therefore removed from the simulated trial after their predicted tumour size reached 2^λ^, where λ denotes the fatal number of tumour doublings for each individual. For each individual, this fatal number λ was obtained by sampling uniformly from {10,11,12,13,14,15}, or the set of possible tumour doubling values between diagnosis and death.

### Optimization Routine for Combined Immuno- and Oncolytic Virotherapies

To provide maximal therapeutic benefit with the lowest possible treatment burden, we defined individualed dosing regimens to be the schedule that minimizes the cumulative tumour burden (the area under the total tumour curve) and the cumulative dose (the total amount of therapy administered over the treatment time). The function to minimize is then expressed as

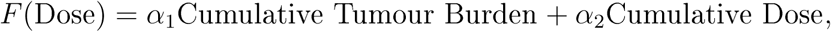

where the positive scaling coefficients *α*_1,2_ weight the importance of maximizing the therapeutic effect versus the need to minimize treatment burden. These weighting values take into account the need for a treatment to be simultaneously effective and tolerable.

Tolerability of combined therapy was attained by bounding the permissible dose size to be four times the standard dose amount. As it is only possible to administer discrete amounts of a drug, typically limited to be some multiple of the available vial size, we constrained the dose size to be 1 – 4-times the standard dose size for both immunotherapy and virotherapy. We allowed for daily immunotherapy dosing and restricted virotherapy administration to days 7, 14, 21, 28, 35, 42, 49, 56, 63, 70 so that virotherapy defined the beginning of a week-long treatment cycle. In total, there are 2000 possible treatment cycles, given that 200 virtual patients underwent ten treatment cycles. Note that the schedule described above is potentially denser than what was administered by Andtbacka et al.^9^ To study whether denser treatment scheduling improved clinical outcomes, we allowed for increased treatment frequency, under the constraint that the cumulative dose administered in the optimal treatment regimen must be less than the cumulative dose administered during the OPTiM trial^9^.

Genetic algorithms are heuristic global optimization routines inspired by natural selection^27-29^ that are frequently employed to estimate parameters in computational biology models. They have also previously been applied to study optimal dosing routines in immunology.^29^ To determine personalized dosing regimens, the optimal function *F*(Dose) was minimized over a ten-week treatment period using Matlab’s genetic algorithm function *ga.*^30^

We then generated personalized schedules for each of the 200 individuals in the optimal combination cohort. These schedules determined an underlying statistical distribution of the likelihood of administering a dose of either immuno- or virotherapy on a given day of the treatment period. Sampling from this empirical distribution, we next determined *P*_*I*_(Day_*i*_) and *P*_*V*_(Day_*i*_), or the probability that immunotherapy or virotherapy, respectively, is administered on Day *i* of therapy. These probabilities served to determine an optimal treatment schedule at the population-level.

### Inference and Validation of Optimal Treatment Schedule

We first, determined whether a dose of immunotherapy was to be administered on the *i*-th day of treatment by flipping a coin based on the probability of administering immunotherapy. This is mathematically equivalent to sampling from a Bernoulli distribution with probability given by *P*_*I*_(Day_*i*_) (see Table 1). If a dose was administered, we sampled from the empirical distribution of dose sizes determined from the individualization (see previous section) with weights P*I*(*n*|Day_*i*_) (i.e. the probability of giving a dose of size *n* given that immunotherapy is administered on day *i)* to determine the size of the immunotherapy size. If the *i*-th day is the beginning of a new treatment cycle, virotherapy may be administered. If so, the same series of steps determined whether virotherapy was administered, and if so, the size of dose, based on *P*_*V*_(Day_*i*_).

**Table 1:** The probability of administering immunotherapy (*P*_*I*_(*A*)) in each day of the treatment cycle and the conditional probability (*P*_*I*_(*n*/*A*) of administering *n* doses of immunotherapy for *n* = 1; 2; 3; 4.

| Day<sub>i</sub> | -3 | -2 | -1 | Start of Cycle | 1 | 2 | 3 |
| --- | --- | --- | --- | --- | --- | --- | --- |
| $P_I(\text{Day}_i)$ | 0.2500 | 0.2580 | 0.2485 | 0.2545 | 0.2465 | 0.2530 | 0.2655 |
| $P_I(1\vert \text{Day}_i)$ | 0.3640 | 0.4109 | 0.3561 | 0.4145 | 0.3753 | 0.3933 | 0.3653 |
| $P_I(2\vert \text{Day}_i)$ | 0.2020 | 0.1802 | 0.1871 | 0.1552 | 0.1866 | 0.1996 | 0.1902 |
| $P_I(3\vert \text{Day}_i)$ | 0.1560 | 0.1376 | 0.1569 | 0.1532 | 0.1562 | 0.1383 | 0.1563 |
| $P_I(4\vert \text{Day}_i)$ | 0.2780 | 0.2713 | 0.2998 | 0.2770 | 0.2819 | 0.2688 | 0.2881 |

To test the effectiveness of the optimal dosing schedule, we created and cloned 100 new virtual patients, and separated them into three three trial arms. The first cohort received the combined immuno- and virotherapy of 125 *µ*g/m^2^ of GM-CSF daily for 14 days in 28 day cycles and 1 dose of virotherapy every 14 days^9^ (standard-of-care). A maintenance therapy schedule was followed for the second cohort (see *Results).* Finally, the dosing regimens determined from the population optimization were applied to the third arm. Mortality and removal from the trial followed the same procedure described in the *Model Calibration* section above.

### Local Sensitivity Analysis

Potential drug targets were identified by performing a sensitivity analysis on the mathematical model’s parameters. Each parameter was varied one-by-one by 10%. The influence of each of these variations was measured by comparing the predicted tumour doubling time and the tumour burden after 15 months to the model’s predictions without any parameter changes according to changes according to

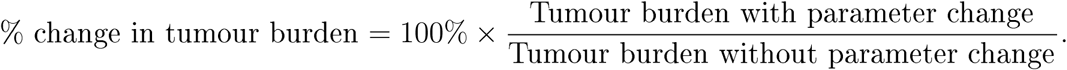

## Results

### Computational Biology Model Successfully Predicts Existing Therapy Results

We first compared the model predictions to the OPTiM results^9^ to evaluate the computational biology model’s ability to accurately represent the outcomes for patients receiving either GM-CSF or the oncolytic virus monotherapy T-VEC.^9,11^ Fig. 3A summarizes the simulated survival fractions. As expected, no untreated virtual patient survived to the end of the trial and both of the treated cohorts display increased survival when compared to no treatment. Patients receiving virotherapy were the most likely to survive until the end of the *in silico* trial.

**Figure 3:**
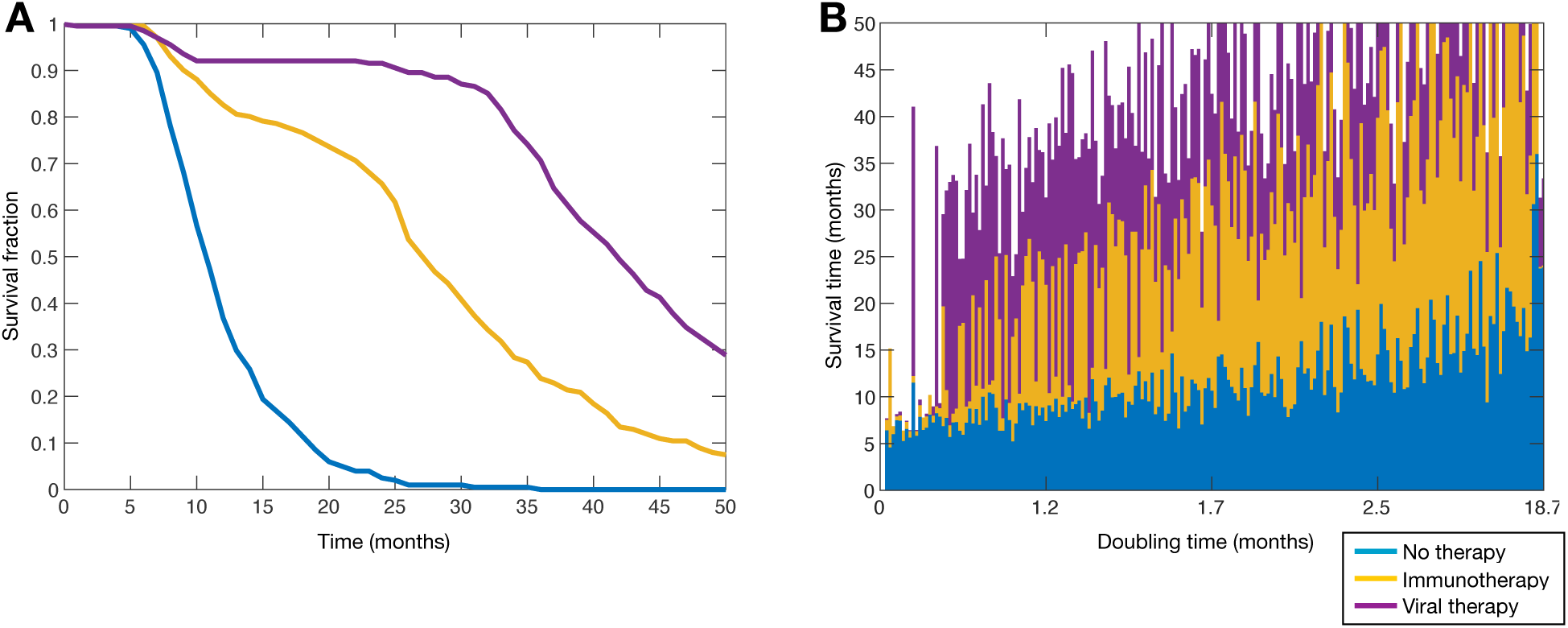
Treatment with oncolytic virus provides improves outcomes over no treatment and immunotherapy in virtual clinical trial. **A)** Kaplan-Meier curves for patients in the no treatment, immunotherapy and virotherapy arms of the virtual trial; **B)** Survival time versus tumour doubling time for virtual patients in each arm of the virtual trial. Overlapping bars represent the outcomes identical triplicates of one virtual patient in each trial arm. Bars that reach the upper limit of Figure (b) represent *in silico* patients who survived to the end of the trial.

The individual beneficial effects of treatment are presented in Fig. 3B. Virtual patients were ordered according to their untreated tumour doubling time, where a longer doubling time indicated slower progression and less aggressive disease. We found that treatment with GM-CSF and oncolytic virus therapy provided larger survival gains in those with with longer doubling times, indicating that oncolytic viruses will have an effect similar to traditional immunotherapy in aggressive cancer strains as evidenced by Figure 4(f) of Andtbacka et al.^9^

**Figure 4:**
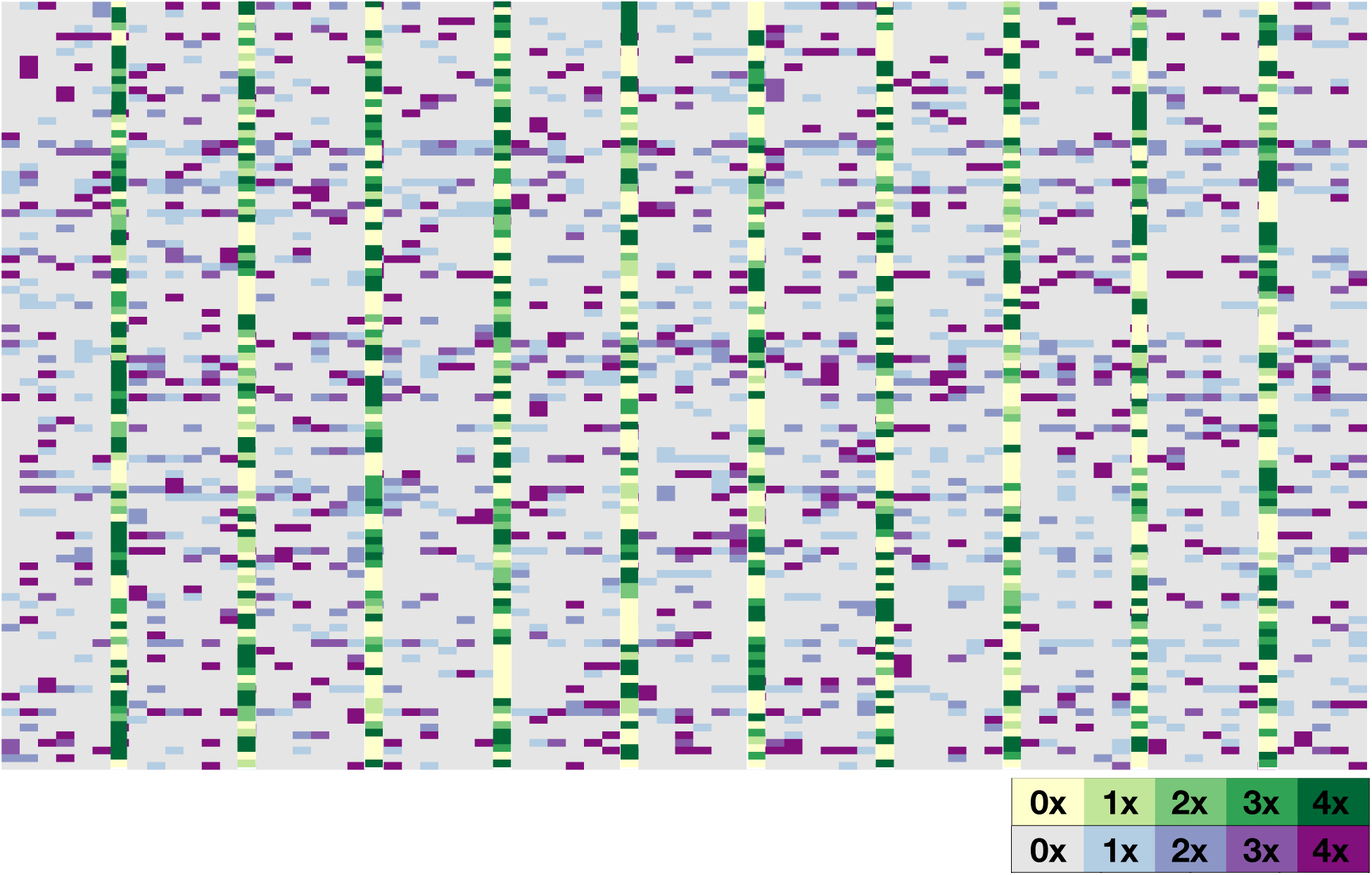
Optimal personalized dose scheduling for each of the 200 virtual patients. The *n*th horizontal row corresponds to the *n*th virtual patient, while the *m*-th vertical column corresponds to the dose administered on day *m*. The dose size is presented as a multiple of the standard dose with immunotherapy in shades of purple and virotherapy in shades of green.

### “All or Nothing” Virotherapy Dosing Strategy

We expected that treatment with GM-CSF would be used to either prime the immune system before virotherapy, or to support the immune response directly following administration of the oncolytic virus. However, as seen in Fig. 4, no structure is easily discerned. To better understand the underlying distribution structure of the individualized treatment schedules, we calculated the probability that any immunotherapy should be administered on each of the seven treatment cycle days of the optimized therapy regimen, as described in the Methods section (*Optimization Routine for Combined Immuno- and Oncolytic Virotherapy).* If a dose was given, we computed the conditional probability of administering a dose of one, two, three or four multiples of the standard dose. We found that the probability of administering a dose of immunotherapy for a given treatment day is roughly constant at 25% throughout the treatment cycle. Interestingly, our results indicate that the immunotherapy dose given is expected to be either the smallest or the largest permitted (see Tab. 1), suggesting that immunotherapy is most useful as an additional instigator when virotherapy does not elicit a sufficient immune response, or to otherwise maintain the immune recruitment initiated by successful viral infection and lysis.

Contrary to the mono-immunotherapy dosing schedule, the conditional probabilities *P*_*V*_(Day_*i*_) in the viral oncology case (see Tab. 2) are heavily skewed to the maximal tolerable dose. Given the mechanism of action of virotherapy (namely, infecting tumour cells), it is unsurprising that administering a larger dose of oncolytic virus should improve clinical outcomes. Put differently, an “all or nothing” approach of dosing infrequently, but for maximal therapeutic benefit, is optimal, in contrast to the logic of the immunotherapy case.

These results suggest that administering immunotherapy between administrations of virotherapy serves mainly to maintain immune recruitment.^31^ To test this hypothesis, we defined Maintenance Therapy to be the administration of virotherapy once every 14 days and with immunotherapy administered evenly throughout on days 3,6,9, and 12 of each virotherapy treatment cycle. Dose size was calculated based on the cumulative expected weekly dose of immunotherapy (8 doses/14 days) from the optimized regimen. Two doses of immunotherapy were therefore administered on days 3,6,9, and 12 to replicate the total expected immunotherapy dose. The same procedure was used to determine the virotherapy dose.

**Table 2:** Conditional probabilities for oncolytic virotherapy administration inferred from individualized dose schedule optimization. The probability of administering virotherapy on each 7th day of the treatment, cycle (*P*_*V*_ (Day_7_)) and the conditional probability *P*_*V*_(*n*|Day_7_) of administering *n* doses of virotherapy for *n* =1,2, 3, 4.

|  |  |
| --- | --- |
| $P_V(\text{Day}_7)$ | **0.5915** |
| $P_V(1\vert \text{Day}_7)$ | 0.1952 |
| $P_V(2\vert \text{Day}_7)$ | 0.1530 |
| $P_V(3\vert \text{Day}_7)$ | 0.1521 |
| $P_V(4\vert \text{Day}_7)$ | 0.4996 |

### Local Sensitivity Establishes Previously-Identified and Potential Therapeutic Targets

The parameters controlling the dynamics of mitotic cells are the most sensitive, with the 10% change accounting for drastic changes in disease burden. For example, increasing the rate at which *G*_1_ cells enter into mitosis results in a 11-fold increase in tumour burden, while decreasing this rate decreases the tumour burden by a factor of 33. This indicates that interventions that inhibit the specific transition from *G*_1_ into mitosis offer therapeutic benefits.

Figure 5 shows the percent change in tumour doubling time and tumour burden corresponding to each parameter.

**Figure 5:**
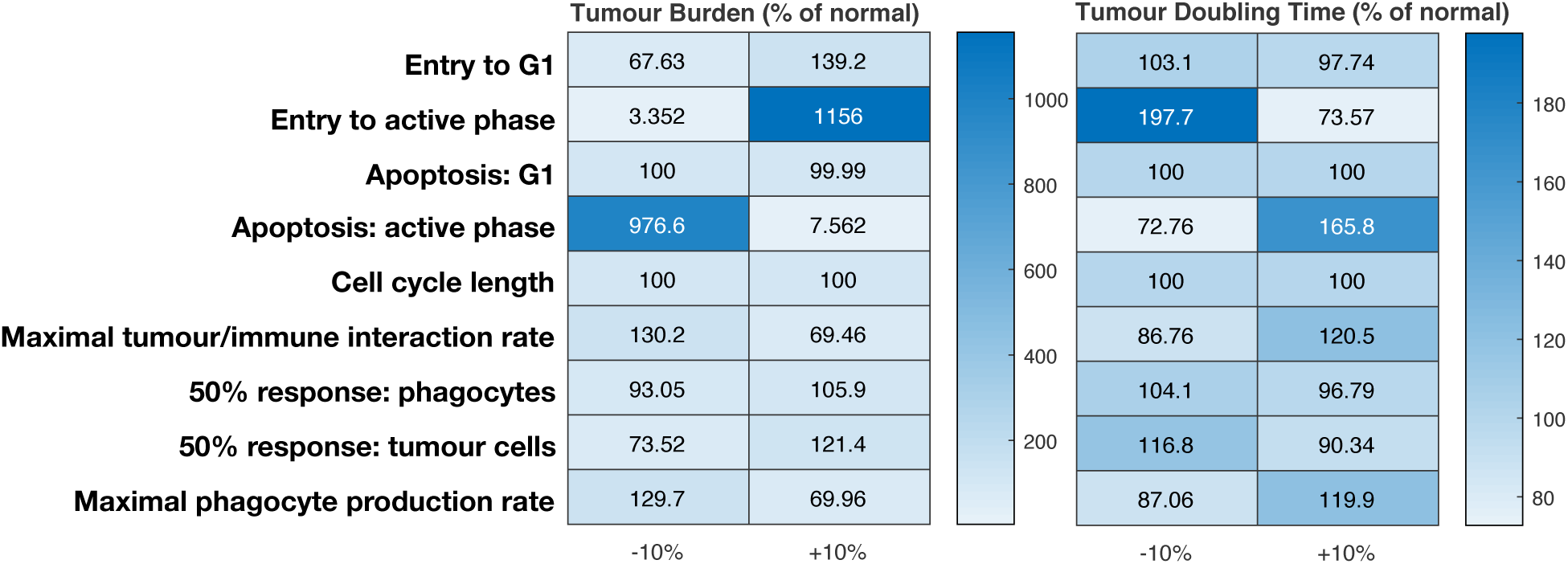
Heat map showing the outcome of local parameter sensitivity analysis. Figure (a) shows the dependence of tumour burden on the parameters shown on the y-axis. Figure (b) shows the dependence of tumour doubling time on the parameters shown on the y-axis. In both cases, parameters were varied by ±10%.

### Maintenance and Predictive Combination Therapies Significantly Improve Patient Survival

Both the standard-of-care (additive combination of the GM-CSF and T-VEC administration protocoles in the OPTiM trial^9^) and optimal combination inimuno- and oncolytic virother-apies improved overall survival times as compared to the OPTiM trials (see Fig. 3). The mean survival time for patients receiving optimized dosing was significantly longer than the mean survival time for patients receiving mono-virotherapy (37.4 months vs. 43.2 months, two-sided t-test p-value of 3.5 x 10^−6^). Maintenance therapy similarly significantly increased mean survival time against mono-virotherapy (37.4 months vs. 42.8 months, two-sided t-test p-value of 1.22 x 10^−5^). The maintenance therapy and optimal dosing regimens also outperformed the standard combination therapy: on average, the mean survival time for patients receiving standard combination therapy was 40.44 months, while patients receiving the maintenance therapy or optimal dosing survived for 42.83 or 43.24 months respectively (two-sided t-test p-values of 0.045 and 0.024). Survival results for the standard and predicted combination therapies results are summarized in Fig. 6.

**Figure 6:**
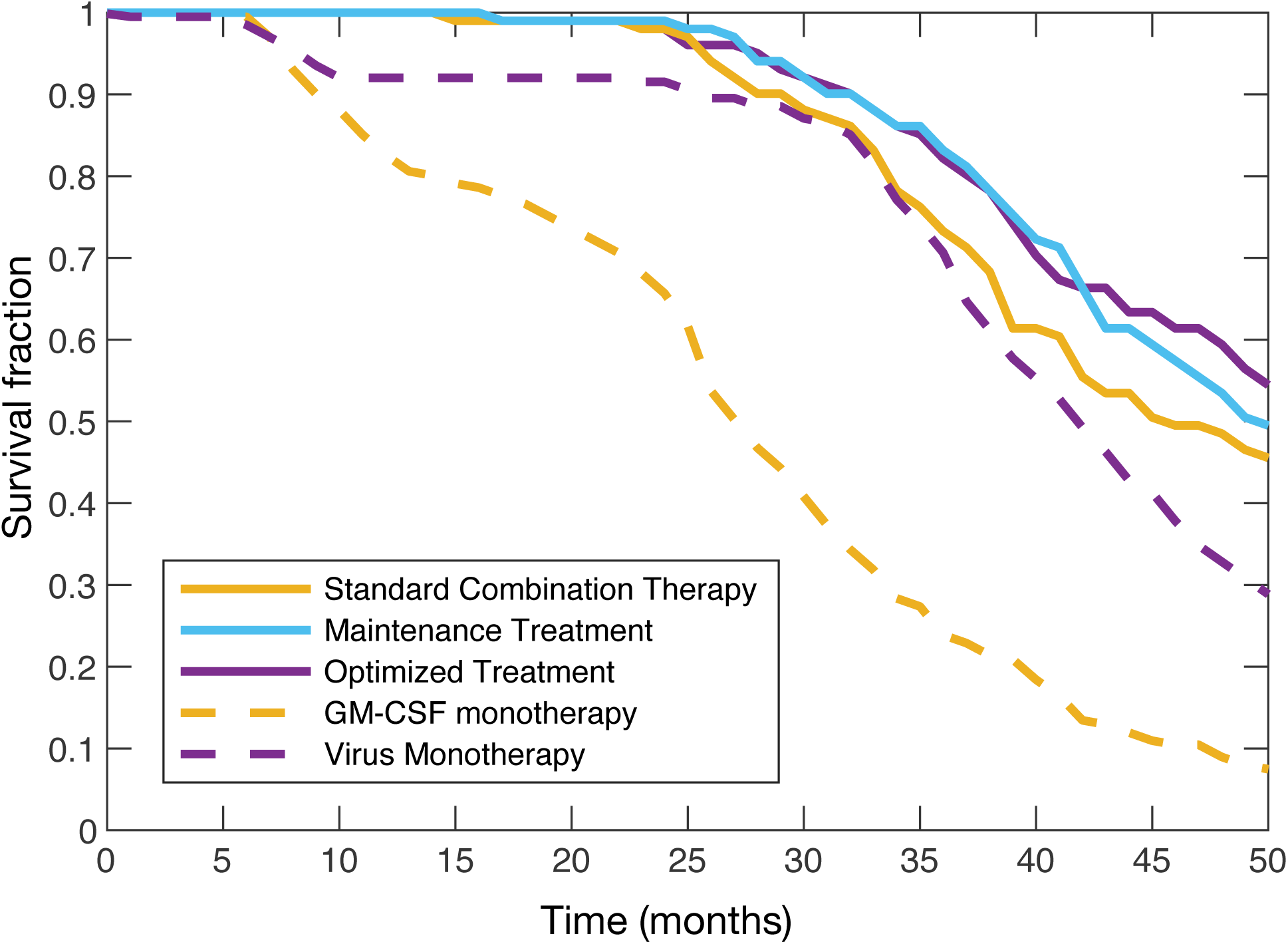
*In silico* clinical trial predicts improved outcomes for both optimal dosing strategies and maintenance therapy versus standard combination therapy. Kaplan-Meier curves for Arm 1: patients receiving Standard Combination Therapy (solid yellow line), Arm 2: Maintenance Treatment (solid cyan), Arm 3: Optimized dosing regimen determined through the *in silico* clinical trial (solid purple), Arm 4: GM-CSF monotherapy (dashed yellow line), and Arm 5: mono-virotherapy (dashed purple). Constrained optimization (Maintenance Therapy) outperforms standard combination therapy, despite additional limitations on dose timing.

To improve therapy tolerability, an additional criterion for regimen optimality is the minimization of the number of treatment days. In the standard of care schedule, patients received 2 administrations of virotherapy and 14 doses of immunotherapy per 28 day cycle, thus requiring 15 total days of drug administration per 28 day cycle. The maintenance therapy schedule required a total of 11 treatment days per 28 day cycle (9 administrations of immunotherapy and 2 administrations of virotherapy), whereas patients given the optimized treatment schedule were administered an expected 7 immunotherapy doses and 2 virotherapy doses per 28 day cycle, for a total of 9 treatment days.

Mean survival times between patients receiving the maintenance therapy and the inferred optimal therapy were not significantly different (42.8 months vs 43.2 months, two-sided t-test p-value of 0.62). This is not unsurprising, given that the maintenance therapy was defined directly from the personalized regimens using the “naïve” constraint that immunotherapy be equally spread out throughout each virotherapy cycle. This restriction was imposed after observing that the probability of administering immunotherapy oil a given day *i* of the treatment cycle (*P*_*I*_ (Day_*i*_)) is more or less equal across our 200 *in silico* individuals. In both cases, the individualization routine was leveraged to create clinically actionable therapeutic strategies. However, as the optimal regimen additionally reduces the overall drug burden during each virotherapy cycle, it could still be considered more optimal than the maintenance strategy, despite the insignificant difference in mean survival time between the two regimens.

In summary, in terms of both end-points and patient burden (defined as the number of treatment days per cycle), immune maintenance therapy outperforms the standard-of-care combination therapy, both of which are improved upon by the optimal dosing schedule. The lack of significant difference between mean survival times in the maintenance and optimal schedules also further motivates the use of *in silico* clinical trial generation as a means to propose candidate investigational regimens prior to clinical trial enrolment.

## Discussion

Improving patient end-points and decreasing the drug burden during anti-cancer treatment are crucial components of cancer care. The introduction of new and advanced therapy modalities is critical to this goal. The approval of T-VEC, the first genetically modified oncolytic virus, was an important step forward for the treatment of late-stage melanoma that significantly improved patient survival over mono-immunotherapy GM-CSF administration. However, the question of whether combined immunotherapy and virotherapy will provide further benefits for patients and, if so, the optimal strategy for such combination therapy, remains. Running clinical trials is an expensive and onerous process. Trial failures are disappointing for patients, clinicians, and researchers, and contribute to overall attrition along the drug development pipeline. Here we have outlined a rational approach to therapy optimization that has significant consequences for how we effectively design and implement clinical trials to maximize their success, and how we treat melanoma, with combined immuno- and virotherapy.

Leveraging our previous computational biology model, we developed an *in silico* clinical trial by creating virtual individuals based on a realistic distribution of model parameter values. Each generated individual was cloned and assigned to different trial cohorts. This innovative strategy enabled us to analyze the effects of distinct therapy procedures on the *same* person, something which is clearly infeasible in the real world. Personalization of treatment regimens was achieved by simultaneously minimizing cumulative tumour and drug burdens. An optimal dosing regimen was subsequently defined based on the resulting personalized treatment schedules. Incorporating clinical realities, we determined that standard combination therapy was improved upon by both a maintenance strategy (where immunotherapy is administered evenly throughout each virotherapy cycle) and this optimal dosing strategy. The latter was shown to be significantly better than both monotherapy and standard-of-care combination approaches.

There are differences between the OPTiM trial and our *in silico* trial. First, without information on the number of individuals in each stage of disease, we cannot identically recreate the underlying distribution of patients in the OPTiM trial. Accordingly, our results are highly dependent on the virtual patients selected for participation based on their tumour doubling time, and would be improved through the incorporation of detailed staging and patient distribution data. Second, the administration of an oncolytic virus can lead to an anti-viral adaptive immune responses and a decrease in treatment efficacy that is currently discounted in the model. Accordingly, the effectiveness of simulated virotherapy does not significantly wane throughout the 12 month treatment period and may account for the observed plateau in survival times during simulated virotherapy. Last, our computational model simplifies tumour-immune interactions by lumping all immune cells into a single phagocyte population. We also considered a single cytokine as a cipher for all pro-inflammatory responses induced by tumour-immune communication. We believe that these considerations do not significantly impact on our general results, but they should be addressed in future work to increase the precision of the predicted personalized regimens. Nonetheless, our results underline the contribution of computational biology to understanding the determinants of improved clinical care and support continued efforts towards rational therapy design. Significantly, this computational biology study suggests promising avenues of investigation towards tailored combination immunotherapy/oncolytic virotherapy for patients with late-stage melanoma.

## Acknowledgements

TC would like t.o thank the Natural Sciences and Engineering Research Council of Canada. (NSERC) for funding through the PGS-D program. MC is grateful for funding through the NSERC Discovery Grant program. Part of this work was performed while TC attended the thematic research program “Mathematical Biology” organized by the Mittag-Leffler Institute. TC and MC would like to thank Tony Humphries for discussions regarding this work.

## Supplementary Information

The computational biology model (described verbally in Eq. (S1)) is the based on that of Cassidy and Humphries^22^ and explicitly includes heterogeneity in tumour cell reproduction velocity and tumour-immune interactions via a distributed delay differential equation. The model describes both quiescent and *G*_1_ phase tumour cell populations while modelling the remainder of mitosis as a delayed process. A phagocyte population and a proinflammatory cytokine, driving the tumour-immune interaction through increased phagocyte recruitment, are also incorporated.

Let *Q*(*t*) and *G*_1_(*t*) denote the quiescent and proliferative phase susceptible tumour cells, respectivcly, and *Q*_*R*_(*t*), *G*_1,*R*_ (*t*) the similar immune-resistant cells. To account for immune selection, we included a resistant, strain of tumour cells that are undetectable to the immune system. We assumed that, tumour cells successfully completing mitosis can randomly mutate into the immune resistant, strain with probability *µ* = 1 x 10^−10^, and we assume that the mutated strain of cancer cells reproduce in an identical manner to the non-mutated strain.

The cytokine concentration is denoted by *C*(*t*), and the phagocyte concentration in the tumour microenvironment by *P(t).* Finally, *V(t)* is the concentration of oncolytic virions and *I(t)* is the number of infected tumour cells. Susceptible tumour cells are infected at a rate *η* while tumour-immune interaction occurs at, the rate *ψ*_*Q,G*_.

Cassidy and Humphries^22^ analysed the mathematical model to determine a threshold tumour-immune interaction to ensure disease regression. Moreover, their analyses showed that cancer progression occurs following a transcritical bifurcation due to decrease immune involvement. This result is consistent with the immunoediting hypothesis.^32^

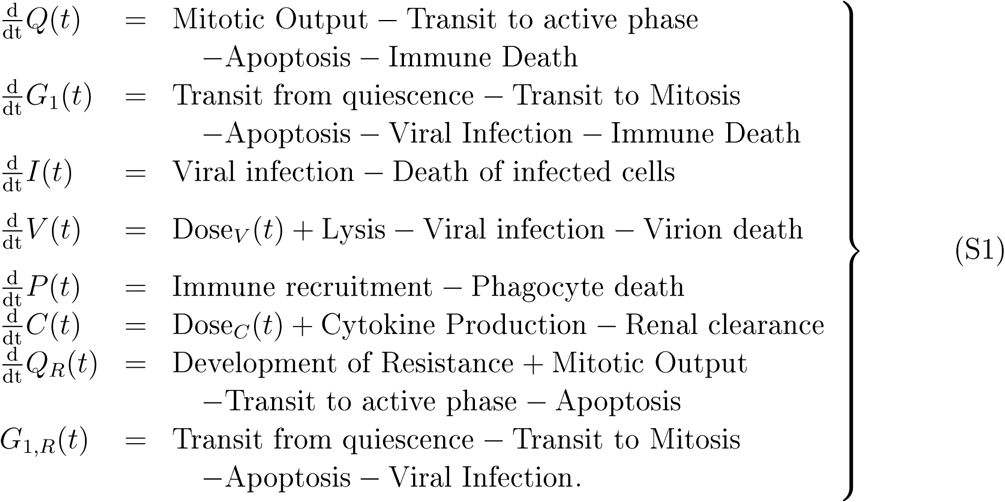

The differential equations describing the progression of disease are

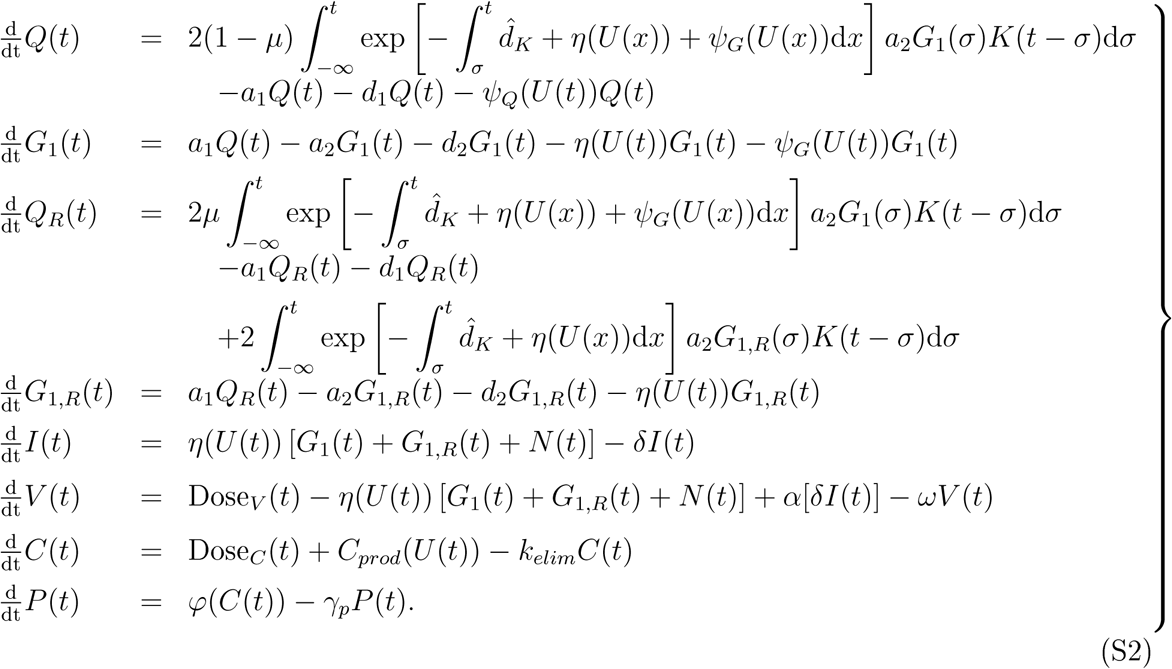

The total number of cells in the cell cycle is given by

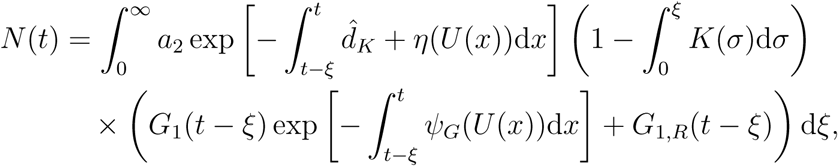

and the parameters used to define the average patient in the section *Generation of In Silico Individuals and Patient Cohorts* in the main text are given in Table S1. These parameters are the same as previously reported.^22^

We model the subcutaneous administration of *N* doses of GM-CSF similar to Craig et al.^33^

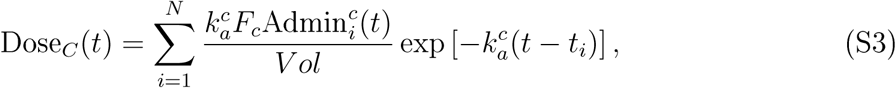

where the amount of GM-CSF administered at time *t*_*i*_ is 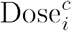 and

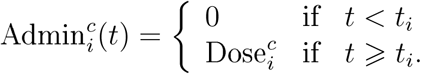

Here, the parameter 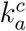 is the absorption rate of GM-CSF, and *F* is the bioavailable fraction of GM-CSF. Similarly, the intralesion administration of oncolytic viruses^9,10,34^ is modelled as

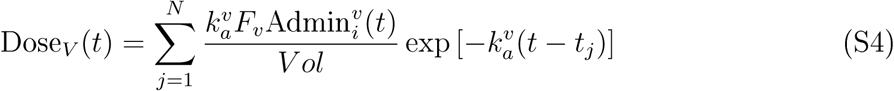

**Table SI:**
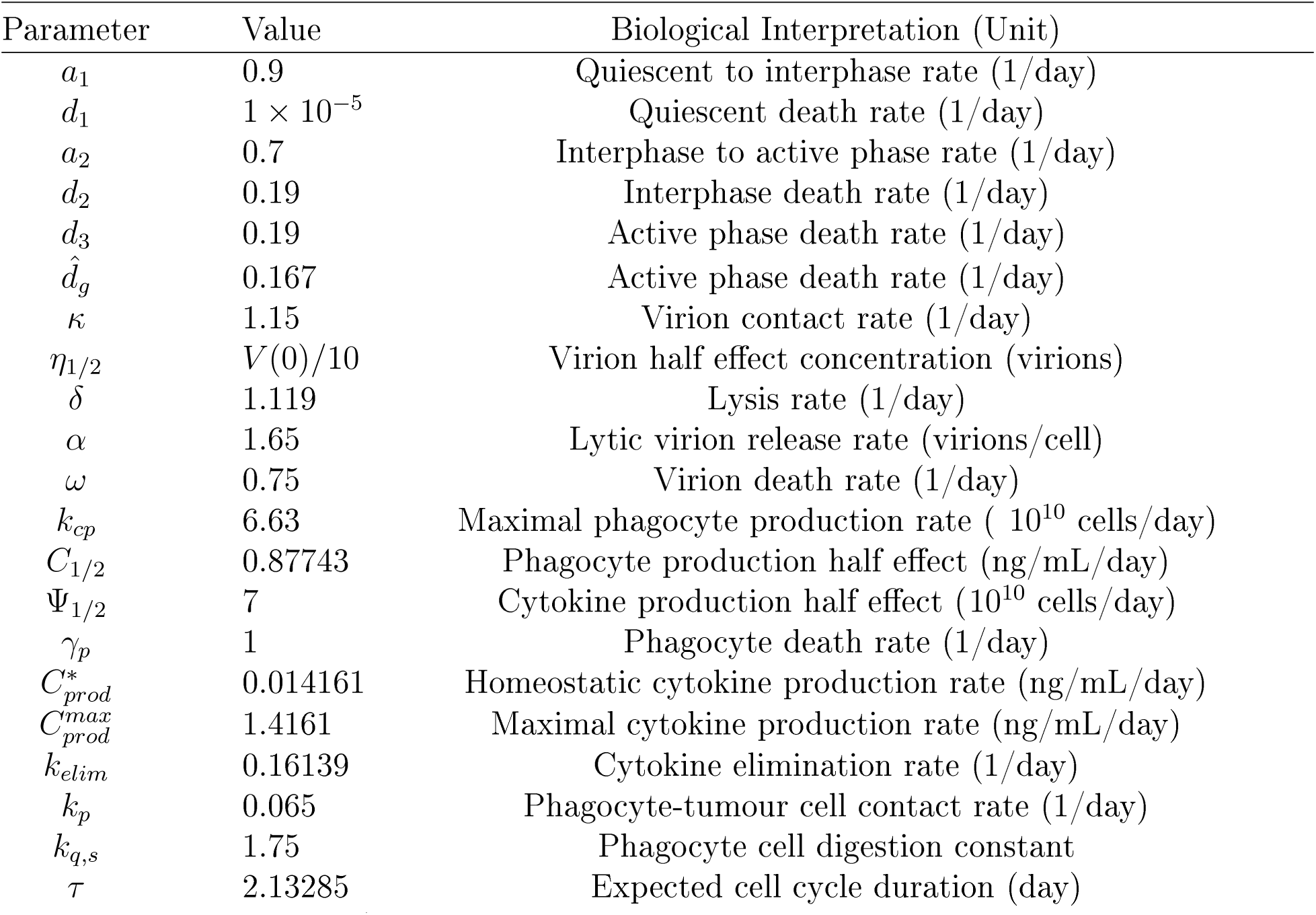
The vector **p** (see section *Generation of in-silico individuals and patient cohorts* in the Main Text.) with biological interpretation. See Cassidy and Humphries^22^ for detailed description of each parameter.

where 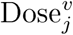 is the amount, of virus administered at time *t = t*_*j*_ and

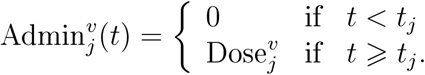

As we consider intralesional administration, the entire dose of oncolytic virus is bioavailable, so *F*_*V*_ = 1 and absorption into the tumour is much faster than cytokine absorption, so 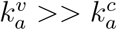. Equation (S2) is simulated using its finite dimensional representation.^22^

### Simulation of the Mathematical Model

Cassidy and Humphries^22^ demonstrated how to reduce the mathematical model (S2) without mutation to a resistant lineage to a finite dimensional system of ordinary differential equations (ODE) when *K(t)* is the probability density function of the gamma distribution,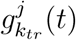. Using an identical technique, the finite dimensional system of ODEs corresponding to (S2) is

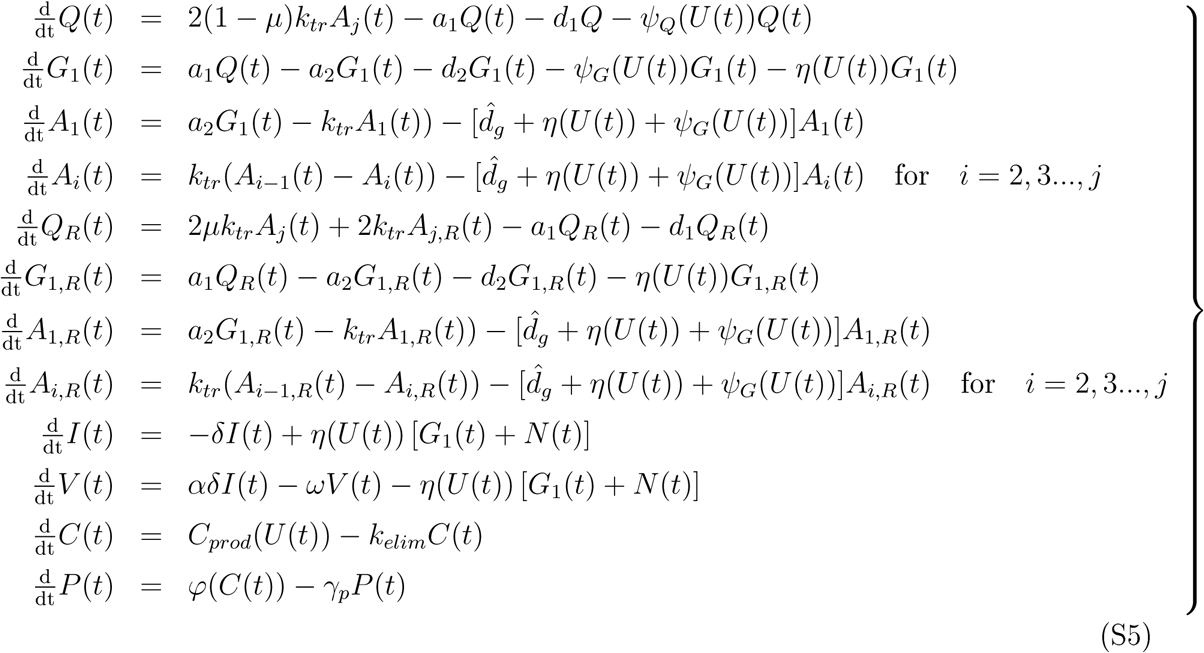

The system in Eq. (S5) is simulated using the stiff ODE solver *ode 15s*.^30^

